# In vitro antimicrobial mechanism of diacerein and potential to reverse MRSE resistance to β-lactams

**DOI:** 10.1101/2023.12.15.571937

**Authors:** Chunyan Fu, Yi Xu, Liping Mao, Chengzhi Zheng, Yangyang Shen, Xinyi Ling, Yumei Zhou, Yiling Yin, Yongliang Lou, Meiqin Zheng

**Affiliations:** National Clinical Research Center for Ocular Diseases, Eye Hospital, Wenzhou Medical University, Wenzhou, China; Eye Hospital and School of Ophthalmology and Optometry, Wenzhou Medical University, Wenzhou; Wenzhou Key Laboratory of Sanitary Microbiology, Key Laboratory of Laboratory Medicine, Ministry of Education, School of Laboratory Medicine and Life Sciences, Wenzhou Medical University, Wenzhou, China

**Keywords:** diacerein, MRSE, antimicrobial mechanism, reversal of drug resistance, β-lactam antibiotics

## Abstract

*Staphylococcus epidermidis* is the most common pathogens causing ocular infection. With the increase of drug resistance rate, it poses a new challenge to anti-infection treatment. In this study, we analyzed the drug resistance of *S. epidermidis* isolated from the eye in the past 5 years to guide empirical antibiotics treatment. Then, the MIC and MBC of diacerein against MRSE were detected, and continuous induced resistance experiments confirmed that MRSE not easily induce resistance to diacerein. In addition, it was observed that diacerein induced MRSE cell lysis, increased membrane permeability and resulted in intracellular ROS accumulation. Diacerein does not have toxic effects to HCEC at effective bacteriostatic concentrations. The results of checkerboard assay indicated that the combination of diacerein and β-lactams had additive effect on MRSE. We also observed that diacerein may reverse MRSE resistance to β-lactam drugs by affecting the active efflux system. In conclusion, our results provide strong evidence for the high therapeutic potential of diacerein against MRSE.

## Introduction

*Staphylococcus epidermidis* that is one of the normal bacterial groups in human skin and mucous membrane is the most common pathogens causing ocular infection^[1]^. Because of the particularity of eye structure, it is necessary to control the infection by empirical antibiotics treatment^[2]^. Unfortunately, in recent years, the problem of drug resistance has become more and more serious, and studies have shown that the drug resistance rate of *Staphylococcus epidermidis* to levofloxacin, the first-line drug for the treatment of eye infections, is increasing year by year^[3]^. In addition, oxacillin-resistant *Staphylococcus epidermidis* (MRSE) has become a major epidemic strain. Although oxacillin is not commonly used to treat eye infections, it has been reported that strains resisted oxacillin can persist in specific environments and promote the spread of drug resistance^[4]^, so the emergence of MRSE poses a new challenge to the clinical anti-infective treatment of ophthalmology. Therefore, it is of great significance to analyze the drug resistance rate find new antibacterial drugs to treat infections caused by MRSE.

There are not enough antimicrobials being developed or used in the clinic. In order to provide more anti-bacterial infection treatment to the clinic, some researchers have concentrated on synthetic materials^[5,6]^, and with the deepening of the concept of repurposing drugs, more and more researchers have focused on drugs that have been used clinically^[7,8]^.

Anthraquinones have attracted more attention because of their remarkable biological activities, such as antivirus^[9]^, anticancer^[10]^, antibiofilm and anti-bacterium^[11]^. The antibacterial activity of diacerein with anthraquinone structure, used for treatment of osteoarthritis^[12]^, has been confirmed by researchers, but its antibacterial mechanism is worth further study^[11,13]^. In the present study, we analyzed the drug resistance of *S. epidermidis* isolated from the Eye Hospital of Wenzhou Medical University in recent 5 years, and confirmed the bacteriostatic activity and explored the antibacterial mechanism of diacerein against MRSE.

## Results

### Analysis of drug resistance characteristics

PBP2a protein expressed by *mecA* gene is the main cause of β-lactam drugs resistance of *S. epidermidis*; therefore, this gene was detected by PCR in 135 strains of *S. epidermidis* isolated from the eyes of patients. The results showed that 81 *S. epidermidis* strains carried *mecA* gene, and the carrying rate was 60%. The results of drug sensitivity analysis showed that the resistance was more than 90% to penicillin, and all of strains were sensitive to vancomycin, amikacin and minocycline. The proportion of multidrug-resistant bacteria which resistant to 3 or more classes of antibiotics in *mecA* strains was 79.12% (64/81), while that in non-*mecA* strains was only 25.92% (14/54), the difference was statistically significant (p < 0.001), so strains carrying the *mecA* gene were more likely to develop resistance to antibiotics commonly used in the clinic. The resistance rates of *mecA* and non-m*ecA* strains were shown in Table 1, and *mecA* strains had higher resistance rates to oxacillin, gentamicin, tobramycin, clindamycin, tetracycline, levofloxacin, norfloxacin, ofloxacin, erythromycin and cotrimoxazole, with statistical significance (p < 0.05).

**Table 1.**
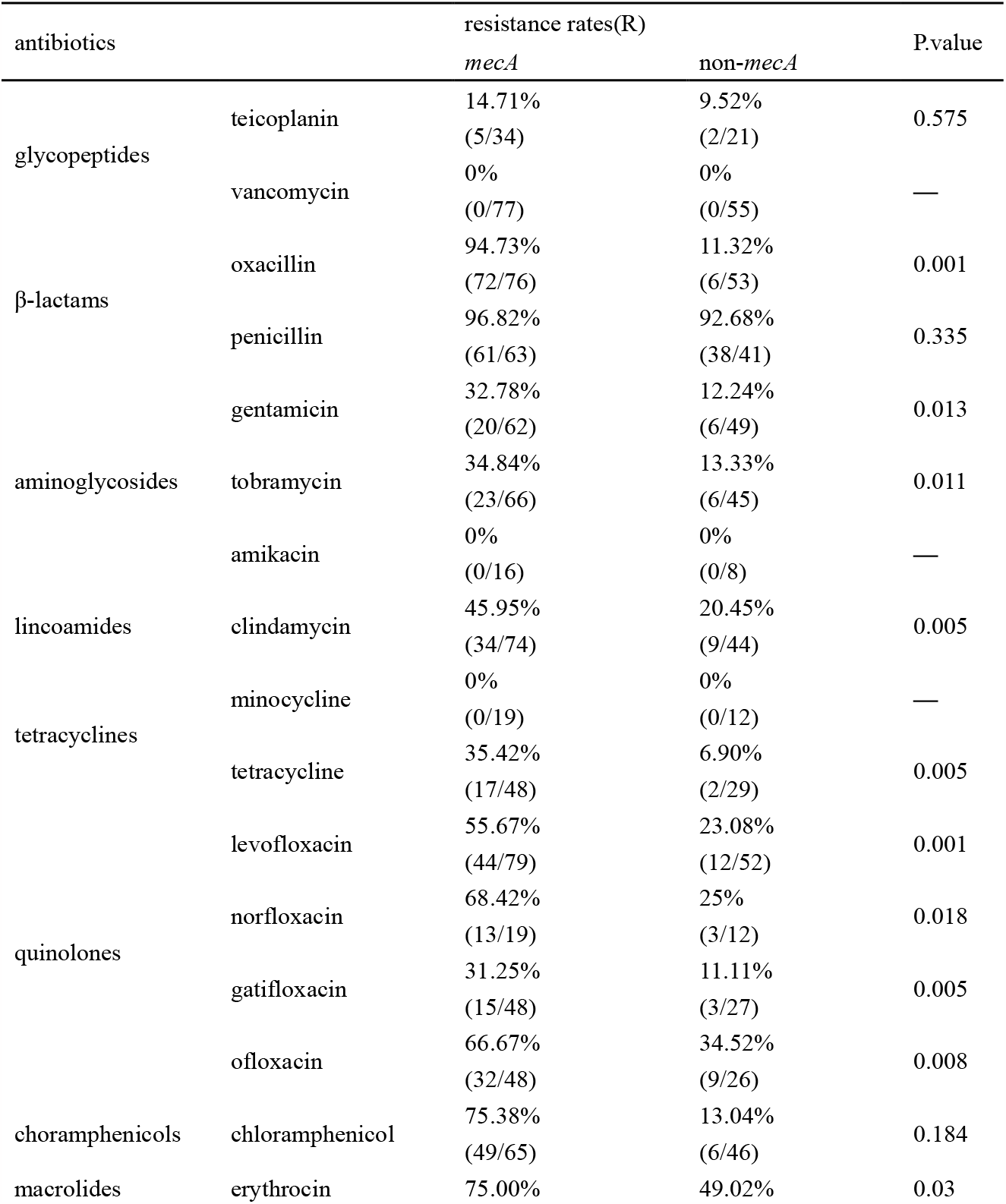
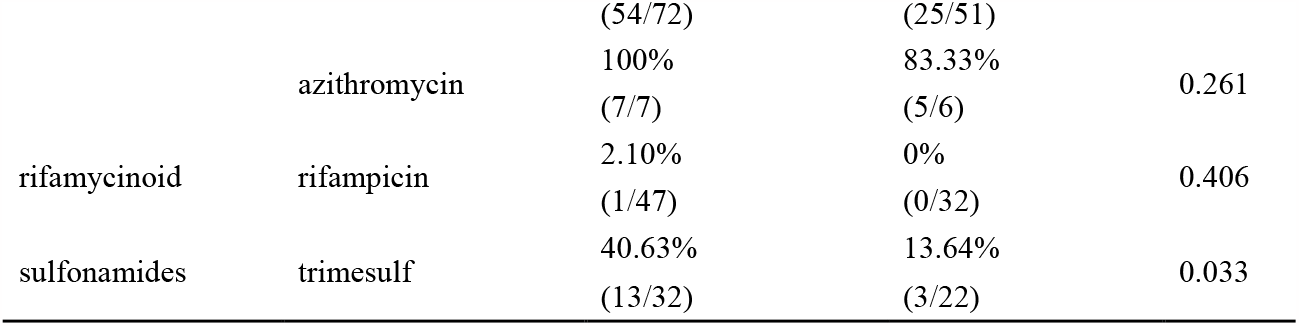
The resistance rates of *mecA* and non-*mecA S. epidermidis* strains to antibiotics commonly used in the clinic.

### Antimicrobial activity of diacerein

Diacerein was tested against 20 clinical strains of *S. epidermidis, S. epidermidis* ATCC 39548 and ATCC12228. The ranges of MIC values and MBC values, modes, and MIC90, MIC50, MBC90, MBC50 are shown in Table 2. MIC90, MBC90, MIC50, MBC50 were the drug concentrations effective against 90% and 50% of strains, respectively.

**Table 2.** MIC and MBC (μg/mL) of diacerein against S. epidermidis.

|  | Phenotypic characteristics | MIC | MIC mode | MIC50 | MIC90 | MBC | MBC mode | MBC50 | MBC90 |
| --- | --- | --- | --- | --- | --- | --- | --- | --- | --- |
| ATCC12228 | Drug-sensitive strains | 4 |  |  |  | 32 |  |  |  |
| ATCC35984 | Resistant to methicillin. *mecA* positive standard strains | 8 |  |  |  | 256 |  |  |  |
| 20 strains of clinical *S. epidermidis* | Resistant to methicillin. *mecA* positive strains | 1-8 | 4 | 4 | 4.4 | 8-256 | 128 | 128 | 256 |

### Cytotoxicity of diacerein in vitro

In order to determine the toxic effect of diacerein on ocular surface cells, CCK-8 and fluorescent staining were used to detect the effect of diacerein on the activity of human corneal epithelial cells HCE-2 (50.B1). In this experiment, the concentration of diacerein was set to be higher than the highest MIC value. The results of CCK-8 experiment showed that only when the concentration of diacerein was more than 20μg/mL, the cell viability was reduced to less than 80% (Table S1).

The results of fluorescence staining showed that when the drug concentration was 15-20μg/mL, the number of cells was less than the control group, indicating that diacerein could inhibit the proliferation of HCEC, but no dead and abnormal cells were found. When the drug concentration was > 30μg/mL, the number of cells with abnormal morphology and dead cells increased significantly, and sheets of cells with circular death appeared (Figure 1). In summary, treatment with MIC concentration of diacerein for 24h did not produce toxic effects to HCEC.

**Fig. 1.**
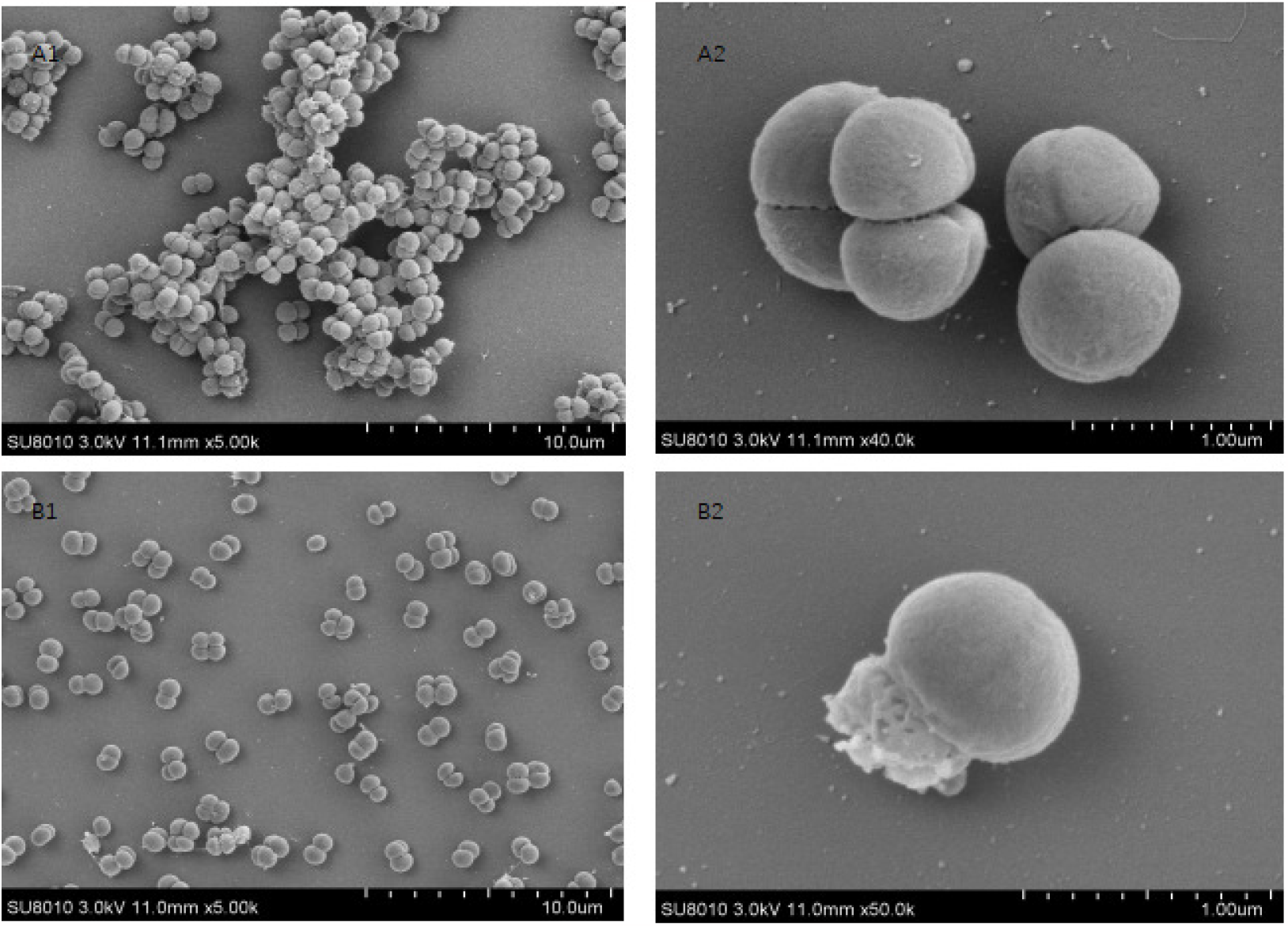
The effect of diacerein on human corneal epithelial cells treated with diacerein for 24h was observed by fluorescence microscope. The green spindle-shaped cells are living cells, the green round cells and the red cells are dead cells.

### Changes in bacterial morphology

For observing the effect of diacerein on the morphology of MRSE, scanning electron microscopy was used to observe the morphological changes of ATCC35984 strain before and after diacerein treatment. As shown in Figure 2, bacteria in the control group grew in clusters, with close links among bacteria and obvious extracellular matrix. The size of the bacteria was uniform, the surface was smooth, and no folds or damage were seen. It suggested that the bacteria had a good shape and could grow and divide normally (Fig.2 A1-2). After treatment with diacerein for 4h, bacterial growth was significantly inhibited, the ability to aggregate and cluster was weakened, the bacteria saw fewer links, the extracellular matrix was rare, most of the bacteria were intact, but the bacteria with damaged morphology were occasionally seen (Fig.2 B1-2). These results indicated that diacerein could inhibit bacterial growth and destroy bacterial structure directly, which led to bacterial cleavage.

**Fig. 2.**
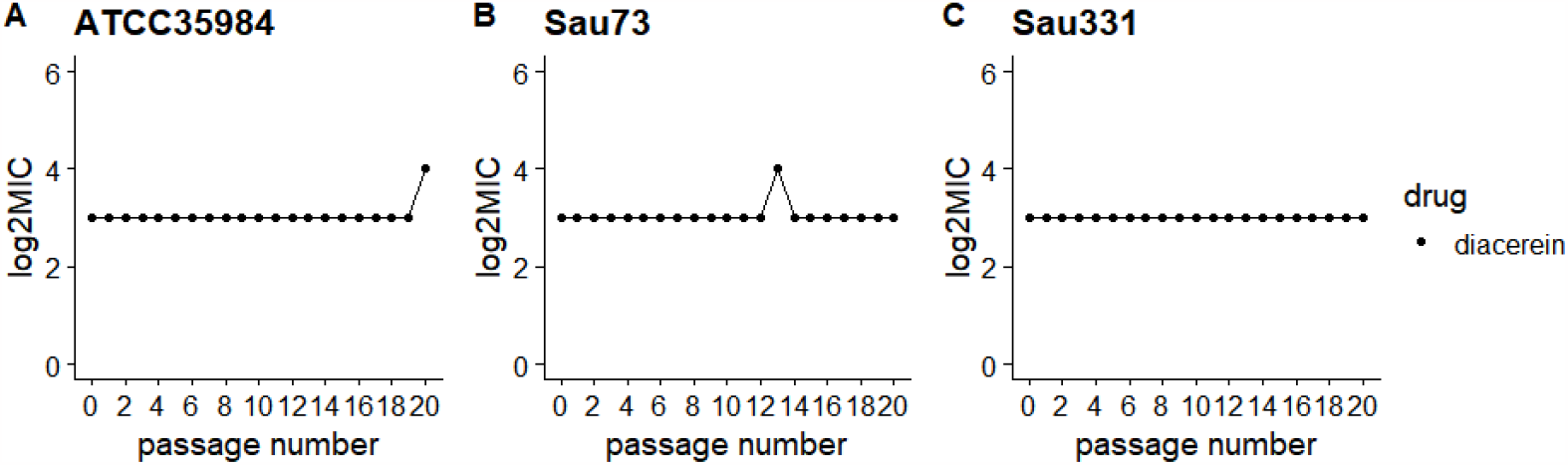
Scanning electron microscopy pictures of *S. epidermidis* ATCC35984 treated with diacerein for 4h. A1-2 and B1-2 respectively show the morphology of bacteria treated with or without diacetylene. A2,B2 has a higher magnification than A2,B1, and the scale is marked at the lower right corner of the pictures.

### Effect of diacerein on the bacterial cell membrane

We tested the effect of diacerein on the bacterial cell membrane permeability of MRSE(ATCC35984, sau73 and sau331) using SYTOX Green probe that can only penetrate into the cell through damaged membranes and fluoresce strongly when it binds to nucleic acids. The intracellular relative fluorescence values of strains (ATCC35984, Sau73 and Sau331) were significantly increased (P < 0.05) under the treatment of low concentration of diacerein, but the values were decreased under the treatment of high concentration (> 2μg/mL) (Fig.3).

**Fig. 3.**
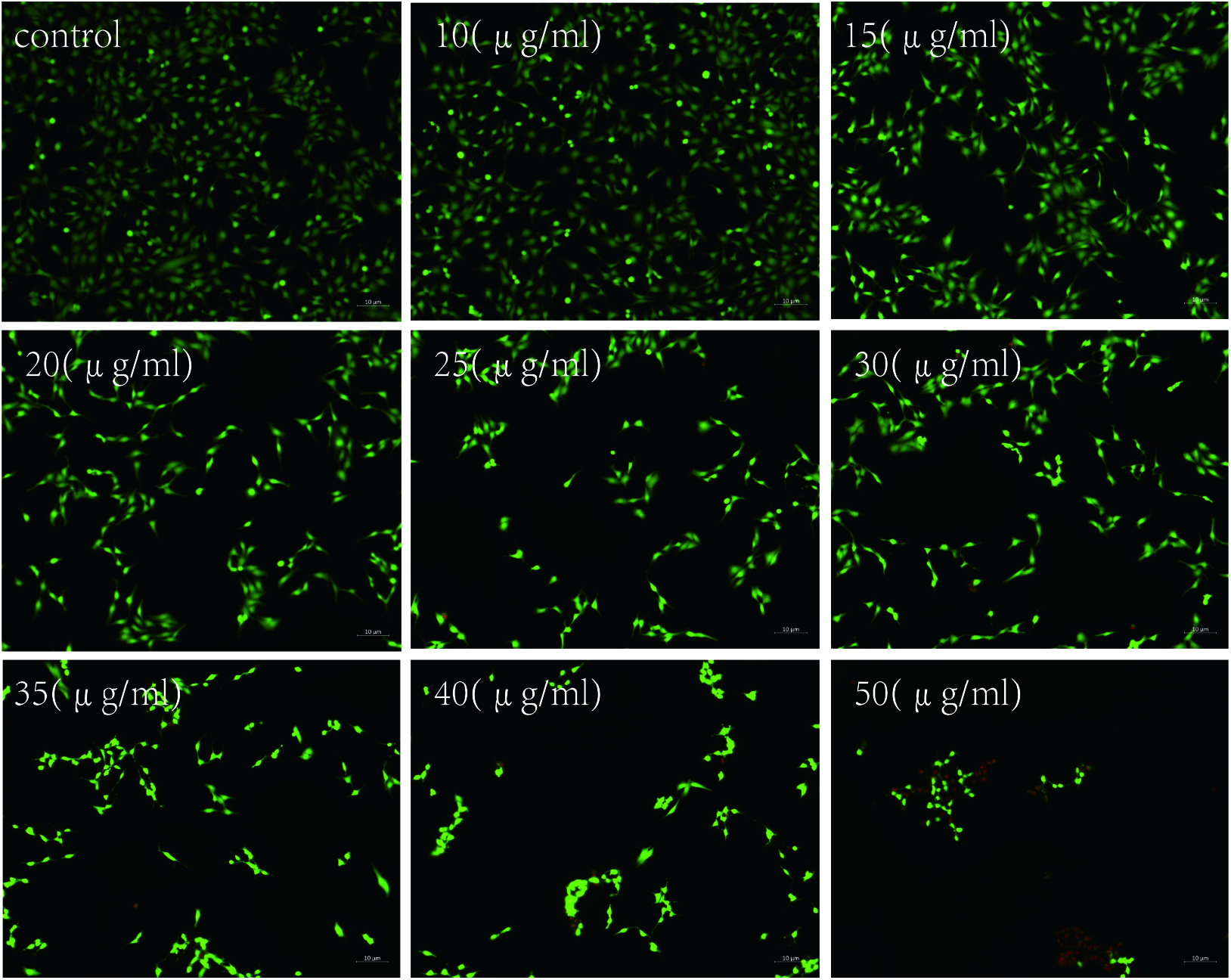
Diacerein increases the permeability of MRSE cell membrane. The group untreated with diacerein was selected as the negative control to calculate the relative fluorescence values, and all values were the mean ±SEM after three repetitions. ***, p<0.001; **,p<0.01; *, p<0.05; ns indicates no significant difference.

In order to explore the reasons why the intracellular relative fluorescence of bacteria under high concentration treatment were reduced, we performed induced autolysis experiments with Triton-X100. Compared with the control group, ATCC35694 strain treated with diacerein had a stronger autolysis capacity, and the difference was most obvious at 1h (Fig.S1). Therefore, the reason for the reduction of intracellular relative fluorescence in MRSE treated with higher concentration of diacerein is that diacerein induces cell autolysis and cell lysis.

### Content of ROS in bacteria

In order to explore whether diacerein affects bacterial growth by increasing ROS accumulation in bacteria, this study used DCFH-DA that can through membrane freely and does not have fluorescence and can be decomposed by intracellular ROS into DCF to detect ROS content in MRSE after treatment with or without diacerein. DCF cannot pass through the cell membrane and has fluorescence, so the level of ROS can be known by detecting the fluorescence of DCF. As shown in Figure 4, the ROS content in the bacteria in the diacerein treatment group was significantly higher than control group, and with the increase of drug concentration, the ROS content also gradually increased. These results indicated that the accumulation of ROS in MRSE strains was in a dose-dependent manner.

**Fig. 4.**
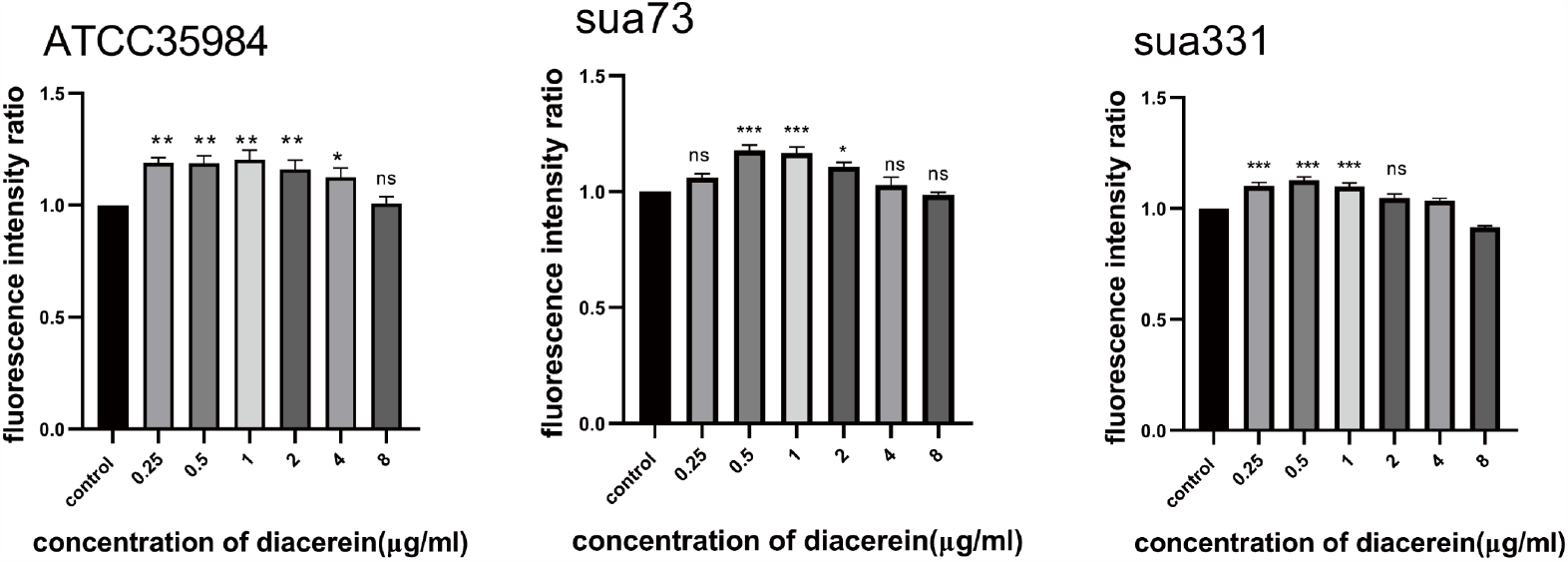
Diacerein can induce accumulate of ROS in MRSE cells. The group untreated with diacerein was selected as the negative control to calculate the relative fluorescence values, and all values were the mean ±SEM after three repetitions. ***, p<0.001; **,p<0.01; *, p<0.05; ns indicates no significant difference.

### Transcriptome enrichment analysis

We explored the effect of diacerein on the gene expression of MRSE(ATCC35984) by transcriptome sequencing and analysis. After differential expression analysis of the data, GSEA analysis based on KEGG database was conducted in this study. The results were shown in Figure S2, four pathways were significantly enriched (p. adjust < 0.05), including three down-regulated pathways (Aminoacyl-tRNA biosynthesis, 2-Oxocarboxylic acid metabolism and valine, leucine and isoleucine biosynthesis) and one up-regulated pathways (homologous recombination).

### Diacerein is not easy to cause MRSE resistance

Induced resistance assays were conducted on strains ATCC35984, Sau73 and Sau331, and changes in MIC values were continuously monitored (Fig.5). The results showed that the MIC value for ATCC35984 changed from 8μg/mL to 16μg/mL only in the 20th generation. The MIC value for strain Sau73 only changed from 8μg/mL to 16μg/mL in the 13th generation, and then dropped back to 8μg/mL in the 14th generation, so the increase in MIC value may be caused by errors. The MIC value for Sau331 strain did not increase in 20 successive generations. A 4-fold increase in MIC value was used as the criterion to judge the strain’s resistance to antibiotics^[14]^, and diacerein was not easy to induce MRSE resistance.

#### Combined drug sensitivity results

Prior research has suggested that diacerein can enhance the antibacterial effect of oxacillin on Staphylococcus aureus in vitro^[15]^. In this study, the combined drug sensitivity test of four β-lactam drugs and diacerein was carried out for MRSE, and FIC values was shown in Table 3. Results indicated that the combination of diacerein and β-lactams had additive effect on MRSE.

**Table 3.** Combined drug sensitivity test of β-lactams and diacerein for MRSE.

| strains | FIC |  |  |  |
| --- | --- | --- | --- | --- |
|  | oxacillin | cefotaxime | imipenem | cefalexin |
| Sau73 | 0.75 | 1 | — | — |
| Sau331 | 0.75 | 1 | — | — |
| ATCC35984 | — | 0.625 | 0.625 | 1 |

### Changes of β-lactam drug sensitivity

Broth microdilution method was used to detect the changes of MIC values of 2 clinically isolated MRSE strains treated with β-lactam drugs before and after 20 generations of continuous diacerein treatment. The results showed that MIC values of oxacillin, cefotaxime and imipenem in strain Sua331 and cefotaxime in strain Sau73 were decreased after treatment with diacerein (Table 4).

**Table 4.** Changes of MIC of β-lactam drug for MRSE.

| strains | MIC(μg/mL) |  |  |
| --- | --- | --- | --- |
|  | oxacillin | cefotaxime | imipenem |
| Sau73-control | 1 | 8 | ≤0.125 |
| Sau73-dia | 1 | 4 | ≤0.125 |
| Sau331-control | 2 | 8 | 0.25 |
| Sau331-dia | 0.5 | 4 | ≤0.125 |

### Possible cause of reversal of MRSE resistance by diacerein

In order to explain the reason for the decreased MIC value of β-lactam drugs, we first ruled out the possibility of bacterial contamination and loss of resistance genes. Whole genome DNA of sau331-control, sua73-control, sau331-dia and sau73-dia strains was extracted, and sequencing and pan-genomics analysis were conducted. The evolutionary tree constructed according to the core genes was shown in Figure S3A. The strains Sau331 and Sau73 treated with or without diacerein were in the same evolutionary branch respectively and had a relatively close kinship, indicating that the cause of MIC value change was not bacterial contamination. We used PCR to detect the *mecA* gene in the genome, and confirmed that the *mecA* gene still existed (Fig. S3B), suggesting that the reason was not losing of *mecA* gene.

Then, we performed transcriptional analysis of sua331-control and sau331-dia strains. A total of 192 differentially expressed genes were identified, including 106 down-regulated genes and 86 up-regulated genes (Fig. S4). The results of GSEA analysis based on KEGG^[16]^ database was shown that 10 pathways were significantly enriched (p. adjust < 0.05) (Fig. 6A-C), consist of four down-regulated pathways (including ABC transports) and six up-regulated pathways. GO functional enrichment was performed (Table S2), and differentially expressed genes were concentrated in phosphoenolpyruvate-dependent sugar phosphotransferase system or essential components of cell membranes. The genes on the bacterial cell membrane are involved in many functions, including cell wall synthesis, efflux pump function, ion transporter, serine protease and so on.

**Fig. 5.**
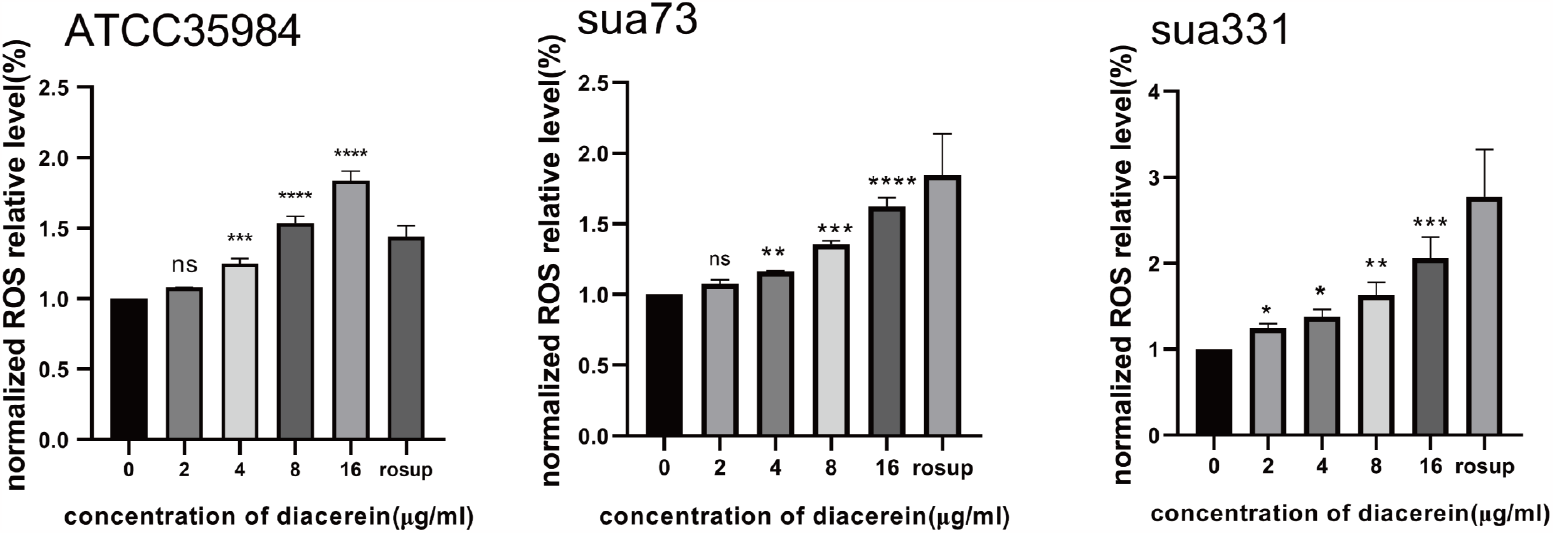
Curves show the continuous change in the MIC value of diacerein against MRSE, which indicated multi-passage resistance selection of MRSE to diacerein.

**Fig. 6.**
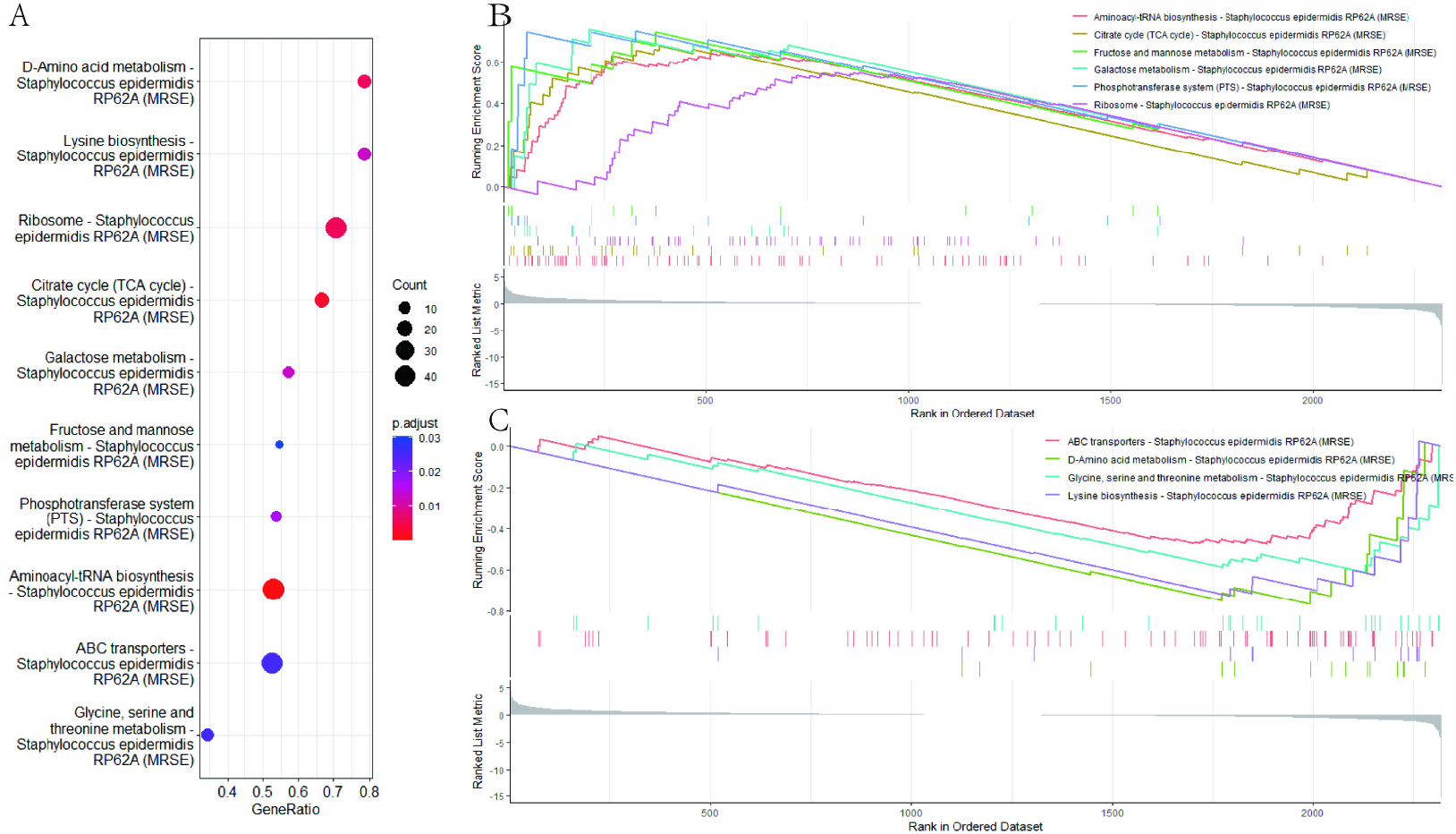
Gene set enrichment analysis. Picture A shows the pathway significantly enriched, the color of the dots represents the p.adjust, and the size of the dots represents the number of genes enriched in the corresponding pathway. B and C shows the downregulated and upregulated pathways, respectively.

## Discussion

*Staphylococcus epidermidis* is the most common pathogenic bacterium causing ocular infection, and the drug resistance rate is increasing year by year, so knowing the drug resistance situation and searching for new antimicrobial agents are important measures to deal with this situation.

As MRSE has become one of the clinically prevalent strains, we analyzed the carrying rate of *mecA* gene and drug resistance of 135 *S. epidermidis* strains isolated from the eyes of patients in the Eye Hospital of Wenzhou Medical University. No strain resistant to vancomycin and amikacin was found, so the current clinical treatment using vancomycin and amikacin to intravitreal injection is still applicable. In addition, it is noteworthy that compared with the strains without *mecA* gene, MRSE strains showed higher resistance to a variety of antibiotics commonly used in clinic, such as β-lactam, tetracyclines, aminoglycosides and chloramphenicol, and the proportion of multi-drug resistant bacteria was also higher. Therefore, we should be vigilant about the prevalence of MRSE strains.

In this study, we confirmed the antibacterial activity of diacerein against MRSE. Due to the MBC to MIC ratio >4, diacerein could be defined as a bacteriostatic drug rather than a bactericidal drug^[17]^, and the result of induced resistance assay showed that diacerein was not easy to induce resistance to MRSE.

Cytotoxicity assay is essential before a drug can be used in human therapy. At present, it has been proved that diacerein has bactericidal effect in mice, and it is not irritating to rabbit eyes^[13]^. As previous study did not conduct experiments on human cells, this study supplemented the cytotoxicity assay of diacerein on HCEC. The results indicated that HCEC in vitro could tolerate diacerein at the concentration higher than 3 times of MIC_90_(4.4μg/mL) for 24h. Therefore, at the concentration inhibiting the growth of MRSE, diacerein did not have toxic effects on HCEC. The serum concentration of most antibacterial drugs should be higher than the MIC at least during the administration interval to achieve clinical antimicrobial efficacy^[18]^. There is no doubt that local treatment could increases the local concentration, so diacerein has the potential to be used as a local treatment for eye infection. However, we need to know the Pharmacokinetic/pharmacodynamic parameters of diacereins in eyes in the future to guide clinical treatment.

Cell membrane is the main barrier that controls the entry and exit of the macromolecules, and is also the synthesis site of bacterial cell wall^[19]^. Therefore, the integrity of cell membrane structure and function is of great significance for the complete life activities of bacteria. In this study, scanning electron microscopy was used to observe that diacerein would cause the cell of MRSE strains to lyse, and diacerein could enhance the bacterial cell membrane permeability by SYTOX Green nucleic acid.

ROS is a natural product of respiratory metabolism, but too much ROS can lead to oxidative damage^[20]^. Inducing excessive ROS accumulation in bacteria is the mechanism by which many antimicrobial agents exert their antibacterial effects^[17]^. We confirmed that diacerein can induce ROS accumulation in bacteria with a dose-dependent manner. The transcriptomic results showed that diacerein can cause protein synthesis to be blocked, but the expression of homologous recombination pathway is upregulated, which may be the result of stress after ROS accumulation to repair genes damaged by ROS.

It was observed that the combination of diacerein and β-lactam drugs had an additive effect on MRSE, and the MIC of β-lactam drugs were decreased after treatment with diacerein. In order to explore the possible causes of the above phenomenon, after ruling out the possibility of contamination and loss of resistance genes, we conducted transcriptome analysis of the strains before and after treatment of diacerein, and the results showed that the expression of genes related to efflux system (including ABC transports) and LCP family proteins was reduced.

Bacterium can expel intracellular drugs through an active efflux system, which results in bacterial drug resistance. Active efflux systems can be divided into several families according to the homology of amino acid sequences, such as ATP-binding cassette (ABC), staphylococcal multidrug resistance family (SMR), resistance-nodulation division family (RND), major facilitator superfamily (MFS), and multi-antimicrobial and toxic compound extrusion (MATE)^[21]^. The complete efflux system consists of interior membrane transporter, membrane channel protein, connexins between the two, and all of which are indispensable and actively pump the intracellular matter out of the cell after proton exchange or hydrolysis of ATP energy. The target of β-lactam drugs is PBPs on the bacterial intima. Due to the poor permeability of the bacterial intima to β-lactams, these tend to accumulate in the Intermembrane space and act on the target proteins. However, previous studies have shown that interior membrane transporter actively capture drugs and exclude them out^[22]^. Therefore, the active efflux system also plays a certain role in the mechanism of bacterial resistance to β-lactam drugs, especially the ABC transports^[23]^.

The cell wall of Gram-positive bacteria contains a variety of sugar-containing polymers, and these substances are often covalently linked to peptidoglycan. LytR-CpsA-Psr (LCP) family proteins^[24]^ and transglycosylase^[25]^ play an important role in the linking of these two substances. Therefore, LCP protein and transglycosylase is indispensable in the synthesis and assembly of bacterial cell membranes and cell walls. Previous studies shown that the loss of LCP protein can damage the cell membrane integrity of streptococcus, group B and make the bacteria sensitive to penicillin^[26]^. In Staphylococcus aureus, the absence of these proteins reduces the bacterial resistance to antibiotics and reduces the ability to infect mouse^[27]^, and some researchers have also regarded LCP protein and transglycosylase as potential targets against drug-resistance^[28]^. In brief, the transcriptome results suggest that diacerein may play a role in reversing MRSE resistance to β -lactam drugs by affecting LCP proteins and the active efflux system. However, the function of LCP proteins and ABC transporters in S. epidermidis resistance to β-lactam drugs has not been clarified, so the above hypothesis also needs knockout or overexpression to verify the function of the gene, which is also the content that needs to improve in the future.

In conclusion, we confirmed the bacteriostatic activity and mechanisms of diacerein against MRSE, the safety to HCEC, and found that it may have the potential to reverse drug resistance. Diacerein become a possible option for the treatment of infections by MRSE.

## Materials and methods

### Bacterial strains and cell line

135 strains of *Staphylococcus epidermidis* were obtained from patients attending the Affiliated Eye Hospital of Wenzhou Medical University, China, between 2016 and December 2022, and all of strains were identified by automatic microbial mass spectrometry (Autof ms600, Autobio, Zhengzhou, China). *S. epidermidis* ATCC 35984 (RP62A), *S. epidermidis* ATCC 12228 are kept in our lab. Human corneal epithelial cells (HCE-2 (50.B1)) were kindly donated by Dr Quan-gui Lin, Affiliated Optometry Hospital of Wenzhou Medical University.

### Detection of *mecA* gene and analysis of antibiotic susceptibilities

Bacterial DNA were extracted by boiling, performing PCR to detect *mecA* gene according to reagent instructions (vazyme, Nanjing, Chain), and primers sequences were as shown in previous^[14]^. *S. epidermidis* ATCC 35984 and 12228 were used as positive and negative controls, respectively. The proportion of strains carrying the *mecA* gene in *S. epidermidis* was counted, then grouping the strains into *mecA* and non-*mecA*, the resistance to clinically used antimicrobial drugs and the multi-resistance rate were counted separately. Chi-square test was performed using SPSS (v26.0) p<0.05 as statistical significance.

### MIC and MBC of diacerein assay

The minimum inhibitory concentrations(MIC) of diacerein against 20 MRSE strains which are carrying the *mecA* gene were assayed in 96-well plates using broth microdilution methods^[13]^. Briefly, diacerein stock solution dissolved in DMSO at a concentration of 20480μg/mL was diluted by MHB and 100μL of working solution was added to 96-well plates containing 100μL of 2 × 10^5^ CFU/mL bacterial suspension, and plates were incubated at 37°C for 24h. MIC is the lowest drug concentration at which no significant bacterial growth is seen, and bacterial growth was be monitored by microplate reader (spectra max 190, molecular devices, USA) at 600 nm. Then, 50μL of bacterial solution suctioning from wells that concentration more than MIC was spread to MH agar plate and colonies were counted after incubation at 37°C for 24 hours. The minimum concentration at which the number of colonies is less than 5 is the MBC, which kills 99.9% of the initial bacteria^[29]^.

### Cell cytotoxicity test

The human corneal epithelial cells (HCE-2 (50.B1)) was cultured in DMEM/f12 medium which was supplemented with 10% fetal bovine serum(FBS) and diacerein in different concentrations, cultured at 37°C,5%CO2 for 24h. Meanwhile, DMSO treatment group was used as control. After incubation, the media containing diacerein were removed, and plates were washed with PBS. The media containing 10% CCK-8(MedChemExpress, New Jersey, USA) was added to wells, and the absorbance value at 450nm was read by microplate reader^[30]^ (spectra max 190, molecular devices, USA), then, calculating the relative fluorescence.To observe changes in HCEC morphology and proliferation, cells were treated with Calcein AM/PI (Beyotime Biotechnology, Shanghai, China) double staining^[31]^ and observed with inverted fluorescence microscope (Carl Zeiss AG, Oberkochen, Germany), after cultured with diacerein.

### Observation the morphology of MRSE

*S. epidermidis* ATCC35984 was incubated with 2MBC of diacerein in 6-well containing cell glass coverslips plate containing at 37°C for 4h, and DMSO treatment group was used as control. After cultivation, the supernatant was removed and cell coverslips were washed by PBS. Then, bacterial morphology was fixed by 2.5% glutaraldehyde at 4°C for overnight and dehydrated with gradient ethanol. Finally, coverslips were sent to Wenzhou Institute, University of Chinese Academy of Science for observation of bacterial morphology using scanning electron microscopy, after coated with layer of gold/palladium^[14]^.

### Detection of ROS

The cells of *S. epidermidis* ATCC35984, sau73 and sau331 loaded DCFH-DA probe (Beyotime Biotechnology, Shanghai, China) were treated with 2-16μg/mL diacerein for 1h at 37°C and the ROSup group was used as positive control, meanwhile, the untreaded group was used as negative control. After incubation, the ROS in bacterial cells were detected with maximum excitation and emission spectra at 488 nm and 525 nm, respectively, on a fluorescence spectrometer (spectra max M5, molecular devices, USA). The relative fluorescence value of each experimental group compared with control group was calculated, and one-way analysis of variance was performed using GraphPad Prism (v8.0) p<0.05 as statistical significance.

### Bacterial membrane permeability assay

The cells of *S. epidermidis* ATCC35984, sau73 and sau331 were treated with 0.25-8μg/mL diacerein working solution which contains 3μM SYTOX Green probe (Invitrogen, Shanghai, China) for 1h at 37°C and untreaded group was used as control. After the culture, fluorescence values of all groups were detected with excitation and emission spectra at 488 nm and 523 nm by fluorescence spectrometer (spectra max M5, molecular devices, USA), then, the relative fluorescence values were calculated as above described, and one-way analysis of variance was performed using GraphPad Prism (v8.0).

### Induced autolysis assay

Combined with 0.2% Triton X-100 to determine whether diacerein promotes the autolysis of MRSE^[32]^. *S. epidermidis* ATCC35984 was treated with 2MIC diacerein working solution which contains 0.2% Triton X-100 (Solarbao, Beijing, China) at 30°C for 220rpm in constant temperature shaker (ONuo Instrument Co., LTD, Tianjin, China). After 0, 0.5, 1, 1.5, 2h of treatment, absorbance value of bacterial suspension was detected by microplate reader and the autolysis curve was drawn^[33]^.

### Transcriptomic Sequencing and Analysis

*S. epidermidis* ATCC35984 was cultured with 4μg/mL diacerein and MHB for 24h at 37°C, respectively, and total RNA extracted using bacterial total RNA extraction kit (TianGen, Beijing, China) were delivered to Peisenol Biotechnology Co., LTD for quality testing and transcriptome sequencing. The resulting clean data were compared with whole genome of *S. epidermidis* ATCC35984 by Hista2. Gene expression in transcripts and differential gene expression were analysed by rsubread tool and DESeq2, respectively. Gene set enrichment analysis (GSEA) basing on Kyoto Encyclopedia of Genes and Genomes (KEGG)was performed.

### Induced resistance assay

Changes in drug resistance were detected by observing successive MIC values. After the first detection of the MIC of diacerein against strains ATCC35984, Sau331 and Sau73, the bacteria in wells with sub-MIC cultured were used for the detection of the next generation of MIC, and this process was repeated 20 times.

### Detection of β-lactams drugs’ MIC against MRSE

MICs of β-lactam drugs (oxacillin, cefotaxime, imipenem) against MRSE that treated with or without diacerein for 20 generation were determined by broth microdilution assay as above description.

### Checkerboard assay

This test was based on the checkerboard assay reported by Andrea Miró-Canturri^[34]^. Diacerein was 2-fold serially diluted along the Y-axis of the 96-well plate, and other β-lactam drugs (oxacillin sodium, cefotaxime, imipenem and cefalexin) were diluted along the X-axis, with the highest concentration of each drug being 2MIC and the lowest concentration being 0. Finally, there were two different concentrations of drugs in each hole of the 96-well plate. Each well plates was added with the same amount of 0.5McF bacterial solution as drug solution, and 96-wells plates were cultured at 37°C for 24h. The fractional inhibitory concentration (FIC) of β-lactam drugs was calculated by dividing the MIC of β-lactams in the presence of diacerein by the MIC of β-lactams alone. The criteria for judging the effect of combined administration were: FIC ≤ 0.5 was considered to have synergistic effect; 0.5<FIC ≤ 1 is additive, 1<FIC ≤ 2 is indifference; FIC>2 indicates antagonism.

### DNA extraction, whole genome sequencing and construction of phylogenetic tree

The whole genome DNA of the sau73 and sau331 strains treated with or without diacerein for 20 generations was extracted and sent to Novogene Co., Ltd. (Beijing, China) for quality testing and sequencing. The resulting data were removed sequencing connectors and control by fastp. Spades and quast were used to complete sequence splicing and detect splicing quality, respectively, and the assembled sequences were used for structural and functional annotation by prokka. Then, roary performs pan-genomic analyses based on core genes and RAxML constructs evolutionary tree using maximum likelihood methods. Finally, iTol that is an online tool was used to visualize and beautify evolutionary trees^[35]^. Meanwhile, the remaining DNA was used to detect whether the mecA gene was missing after 20 successive generations by PCR, as what mentioned before.

### Transcriptome analysis of possible causes of reversal of resistance

Total RNA was extracted from strains of Sau331-control and sau331-dia that was y treated with diacerein for 20 generations, and sent to Shanghai Piceno Biotechnology Co., Ltd. for quality detection and prokaryotic transcriptomic sequencing. The basic analysis process of the returned sequencing data was consistent with mentioned above. However, the expression matrix is synthesized with featureCounts, and after the differential expression analysis was performed with DESeq2 software, the genes satisfying the conditions of log2FlodChange > 1 and padj < 0.05 were selected as differential genes. The differential gene volcano map was drawn with R software. Finally, functional annotation and enrichment of differential genes were performed using the Database for Annotation, Visualization and Integrated Discover (DAVID). Finally, GSEA basing on KEGG was performed.

## Acknowledgments

The raw data sets of WGS and transcriptome sequencing in this study have been uploaded to the NCBI database under the accession number PRJNA1050652.

There is no competition of interest in this article and this subject has passed the ethical approval of eye hospital of Wenzhou Medical University, and the approval number is No. 51 of 2023 in eye hospital of Wenzhou Medical University.

Chunyan Fu: designed and performed the experiments, participated in data analysis and wirting manuscript. Yi Xu was the main experiment implementer, too. Liping Mao and Chengzhi Zheng isolated and preserved bacterial strains; Yangyang Shen and Xinyi Ling analyzed the sequencing data; Yumei Zhou and Yiling Yin: participated in data analysis. Yongliang Lou: corresponding author, was be responsible for data curation and writing the manuscript; Meiqin Zheng: corresponding author, was be responsible for supervision, funding acquisition, and reviewed the manuscript. All authors have seen and approved the final version of the manuscript being submitted

## References

[1] Callewaert C, Ravard Helffer K, Lebaron P. Skin Microbiome and its Interplay with the Environment[J]. Am J Clin Dermatol, 2020, 21(Suppl 1): 4–11.

[2] Das T. Endophthalmitis Management: Stain-Culture, Empirical Treatment, and Beyond[J]. Asia Pac J Ophthalmol (Phila), 2020, 9(1): 1–3.

[3] Kang J Y, Lee W, Noh G M, et al. Fluoroquinolone resistance of Staphylococcus epidermidis isolated from healthy conjunctiva and analysis of their mutations in quinolone-resistance determining region[J]. Antimicrob Resist Infect Control, 2020, 9(1): 177.

[4] Jang S. Multidrug efflux pumps in Staphylococcus aureus and their clinical implications[J]. J Microbiol, 2016, 54(1): 1–8.

[5] Wu H, Wei M, Hu S, et al. A Photomodulable Bacteriophage-Spike Nanozyme Enables Dually Enhanced Biofilm Penetration and Bacterial Capture for Photothermal-Boosted Catalytic Therapy of MRSA Infections[J]. Adv Sci (Weinh), 2023, 10(24): e2301694.

[6] Zhang Y, Yue T, Gu W, et al. pH-responsive hierarchical H(2)S-releasing nano-disinfectant with deep-penetrating and anti-inflammatory properties for synergistically enhanced eradication of bacterial biofilms and wound infection[J]. J Nanobiotechnology, 2022, 20(1): 55.

[7] Wang C H, Hsieh Y H, Powers Z M, et al. Defeating Antibiotic-Resistant Bacteria: Exploring Alternative Therapies for a Post-Antibiotic Era[J]. Int J Mol Sci, 2020, 21(3).

[8] Tarín-Pelló A, Suay-García B, Pérez-Gracia M T. Antibiotic resistant bacteria: current situation and treatment options to accelerate the development of a new antimicrobial arsenal[J]. Expert Rev Anti Infect Ther, 2022, 20(8): 1095–1108.

[9] Parvez M K, Al-Dosari M S, Alam P, et al. The anti-hepatitis B virus therapeutic potential of anthraquinones derived from Aloe vera[J]. Phytother Res, 2019, 33(11): 2960–2970.

[10] Nowak-Perlak M, Ziółkowski P, Woźniak M. A promising natural anthraquinones mediated by photodynamic therapy for anti-cancer therapy[J]. Phytomedicine, 2023, 119: 155035.

[11] Fu C, Xu Y, Zheng H, et al. In vitro antibiofilm and bacteriostatic activity of diacerein against Enterococcus faecalis[J]. AMB Express, 2023, 13(1): 85.

[12] Panova E, Jones G. Benefit-risk assessment of diacerein in the treatment of osteoarthritis[J]. Drug Saf, 2015, 38(3): 245–52.

[13] Zhang H, Liu S, Yue J, et al. In Vitro Antimicrobial Activity of Diacerein on 76 Isolates of Gram-Positive Cocci from Bacterial Keratitis Patients and In Vivo Study of Diacerein Eye Drops on Staphylococcus aureus Keratitis in Mice[J]. Antimicrob Agents Chemother, 2019, 63(4).

[14] Liu M, Peng W, Qin R, et al. The direct anti-MRSA effect of emodin via damaging cell membrane[J]. Appl Microbiol Biotechnol, 2015, 99(18): 7699–709.

[15] Nguon S, Novy P, Kokoska L. Potentiation of the in vitro antistaphylococcal effect of oxacillin and tetracycline by the anti-inflammatory drug diacetyl rhein[J]. Chemotherapy, 2013, 59(6): 447–52.

[16] Subramanian A, Tamayo P, Mootha V K, et al. Gene set enrichment analysis: a knowledge-based approach for interpreting genome-wide expression profiles[J]. Proc Natl Acad Sci U S A, 2005, 102(43): 15545–50.

[17] Li T, Lu Y, Zhang H, et al. Antibacterial Activity and Membrane-Targeting Mechanism of Aloe-Emodin Against Staphylococcus epidermidis[J]. Front Microbiol, 2021, 12: 621866.

[18] Craig W A. Does the dose matter?[J]. Clin Infect Dis, 2001, 33 Suppl 3: S233–7.

[19] Ganesan N, Mishra B, Felix L, et al. Antimicrobial Peptides and Small Molecules Targeting the Cell Membrane of Staphylococcus aureus[J]. Microbiol Mol Biol Rev, 2023, 87(2): e0003722.

[20] Mazur P, Skiba-Kurek I, Mrowiec P, et al. Synergistic ROS-Associated Antimicrobial Activity of Silver Nanoparticles and Gentamicin Against Staphylococcus epidermidis[J]. Int J Nanomedicine, 2020, 15: 3551–3562.

[21] Schindler B D, Kaatz G W. Multidrug efflux pumps of Gram-positive bacteria[J]. Drug Resist Updat, 2016, 27: 1–13.

[22] Oliveira-Tintino C D M, Tintino S R, Justino De Araújo A C, et al. Efflux Pump (QacA, QacB, and QacC) and β-Lactamase Inhibitors? An Evaluation of 1,8-Naphthyridines against Staphylococcus aureus Strains[J]. Molecules, 2023, 28(4).

[23] Villet R A, Truong-Bolduc Q C, Wang Y, et al. Regulation of expression of abcA and its response to environmental conditions[J]. J Bacteriol, 2014, 196(8): 1532–9.

[24] Stefanović C, Hager F F, Schäffer C. LytR-CpsA-Psr Glycopolymer Transferases: Essential Bricks in Gram-Positive Bacterial Cell Wall Assembly[J]. Int J Mol Sci, 2021, 22(2).

[25] Guo Y, Pfahler N M, Völpel S L, et al. Cell wall glycosylation in Staphylococcus aureus: targeting the tar glycosyltransferases[J]. Curr Opin Struct Biol, 2021, 68: 166–174.

[26] Rajaei A, Rowe H M, Neely M N. The LCP Family Protein, Psr, Is Required for Cell Wall Integrity and Virulence in Streptococcus agalactiae[J]. Microorganisms, 2022, 10(2).

[27] Li F, Zhai D, Wu Z, et al. Impairment of the Cell Wall Ligase, LytR-CpsA-Psr Protein (LcpC), in Methicillin Resistant Staphylococcus aureus Reduces Its Resistance to Antibiotics and Infection in a Mouse Model of Sepsis[J]. Front Microbiol, 2020, 11: 557.

[28] Huang C Y, Shih H W, Lin L Y, et al. Crystal structure of Staphylococcus aureus transglycosylase in complex with a lipid II analog and elucidation of peptidoglycan synthesis mechanism[J]. Proc Natl Acad Sci U S A, 2012, 109(17): 6496–501.

[29] Rodríguez-Melcón C, Alonso-Calleja C, García-Fernández C, et al. Minimum Inhibitory Concentration (MIC) and Minimum Bactericidal Concentration (MBC) for Twelve Antimicrobials (Biocides and Antibiotics) in Eight Strains of Listeria monocytogenes[J]. Biology (Basel), 2021, 11(1).

[30] Fan Q, Wu H, Kong Q. Superhydrophilic PLGA-Graft-PVP/PC Nanofiber Membranes for the Prevention of Epidural Adhesion[J]. Int J Nanomedicine, 2022, 17: 1423–1435.

[31] Wang S, Xiong Y, Lalevée J, et al. Biocompatibility and cytotoxicity of novel photoinitiator π-conjugated dithienophosphole derivatives and their triggered polymers[J]. Toxicol In Vitro, 2020, 63: 104720.

[32] Bao Y, Zhang X, Jiang Q, et al. Pfs promotes autolysis-dependent release of eDNA and biofilm formation in Staphylococcus aureus[J]. Med Microbiol Immunol, 2015, 204(2): 215–26.

[33] Si W, Wang L, Usongo V, et al. Colistin Induces S. aureus Susceptibility to Bacitracin[J]. Front Microbiol, 2018, 9: 2805.

[34] Miró-Canturri A, Vila-Domínguez A, Caretero-Ledesma M, et al. Repurposing of the Tamoxifen Metabolites to Treat Methicillin-Resistant Staphylococcus epidermidis and Vancomycin-Resistant Enterococcus faecalis Infections[J]. Microbiol Spectr, 2021, 9(2): e0040321.

[35] Lin S, Sun B, Shi X, et al. Comparative Genomic and Pan-Genomic Characterization of Staphylococcus epidermidis From Different Sources Unveils the Molecular Basis and Potential Biomarkers of Pathogenic Strains[J]. Front Microbiol, 2021, 12: 770191.

